# In-field detection and quantification of Septoria tritici blotch in diverse wheat germplasm using spectral-temporal features

**DOI:** 10.1101/664011

**Authors:** Jonas Anderegg, Andreas Hund, Petteri Karisto, Alexey Mikaberidze

## Abstract

Hyperspectral remote sensing holds the potential to detect and quantify crop diseases in a rapid and non-invasive manner. Such tools could greatly benefit resistance breeding, but their adoption is hampered by i) a lack of specificity to disease-related effects and ii) insufficient robustness to variation in reflectance caused by genotypic diversity and varying environmental conditions, which are fundamental elements of resistance breeding.

We hypothesized that relying exclusively on temporal changes in canopy reflectance during pathogenesis may allow to specifically detect and quantify crop diseases whilst minimizing the confounding effects of genotype and environment. To test this hypothesis, we collected time-resolved canopy hyperspectral reflectance data for 18 diverse genotypes on infected and disease-free plots and engineered spectral-temporal features representing this hypothesis.

Our results confirm the lack of specificity and robustness of disease assessments based on reflectance spectra at individual time points. We show that changes in spectral reflectance over time are indicative of the presence and severity of septoria tritici blotch (STB) infections. Furthermore, the proposed time-integrated approach facilitated the delineation of disease from physiological senescence, which is pivotal for efficient selection of STB-resistant material under field conditions. A validation of models based on spectral-temporal features on a diverse panel of >300 wheat genotypes offered evidence for the robustness of the proposed method. This study demonstrates the potential of time-resolved canopy reflectance measurements for robust assessments of foliar diseases in the context of resistance breeding.

## 1 Introduction

Hyperspectral remote sensing has shown significant potential for the rapid, non-invasive assessment of crop diseases at different scales, ranging from single leaves (*e.g.* Ashourloo et al., 2014; Mahlein et al., 2010) to the canopy (*e.g.* Cao et al., 2013; Yu et al., 2018) to fields and regions (Wakie et al., 2016). Applications have been proposed primarily in the context of precision agriculture, but resistance breeding may equally benefit (Mahlein, 2016). The identification of novel sources of durable, quantitative disease resistance requires screening large and diverse germplasm collections under field conditions. Reflectance-based techniques hold the potential to reduce associated costs and allow for the screening of more genetic variation, if deployed on adequate phenotyping platforms (Aasen et al., 2018; Aasen and Bolten, 2018; Kirchgessner et al., 2017). This may enable indirect selection in early breeding generations and facilitate the identification of novel sources of resistance.

However, to benefit crop breeding, new methods must accurately estimate phenotypes for large numbers of diverse genotypes under field conditions (Araus et al., 2018; Araus and Cairns, 2014; Furbank and Tester, 2011). This represents a significant challenge because genotypic diversity and contrasting environmental conditions are major sources of variation in spectral reflectance. This variation arises mostly from (i) genotype morphology, canopy cover and canopy 3-D structure (Gutierrez et al., 2015; Haboudane et al., 2002; Jacquemoud et al., 2009; Zarco-Tejada et al., 2005), (ii) differences in genotype phenology and the timing of developmental transitions such as heading, flowering and senescence (Kipp et al., 2014; Pimstein et al., 2009; Stuckens et al., 2011) and (iii) reactions to other biotic or abiotic stresses, which may result in similar spectral responses as the disease of interest (Zhang et al., 2012). At present, effects of diseases on canopy reflectance are often investigated at specific points in time (*see e.g.* Cao et al., 2013; Yang, 2010; Yu et al., 2018). Such investigations have provided valuable but highly context-specific insights (i.e. specific to the genotype, growth stage and/or site and environment under study; *see e.g.* Delalieux et al., 2007; Zhang et al., 2012; Zheng et al., 2019). Accordingly, identified spectral features and corresponding thresholds or calibration curves are not sufficiently robust (i.e. universally applicable) for use in resistance breeding. Largely due to such difficulties, high throughput phenotyping of disease resistance under field conditions using hyperspectral reflectance is still elusive (Araus et al., 2018).

Septoria tritici blotch (STB) caused by the fungal pathogen *Zymoseptoria tritici* is a serious threat to wheat production in major wheat growing areas around the world (Orton et al., 2011; Torriani et al., 2015). The development of cultivars with improved resistance to this disease has become a significant objective in wheat breeding and constitutes a key component of STB management strategies (Brown et al., 2015; McDonald and Mundt, 2016; O’Driscoll et al., 2014). Several major resistance genes conferring near-complete resistance to certain *Z. tritici* isolates have been identified and used in commercial cultivars (*reviewed by* Brown et al., 2015). However, these genes are frequently overcome within a few years of their introduction due to the high evolutionary potential of *Z. tritici* populations (McDonald and Mundt, 2016). Genetic loci conferring broadly effective partial resistance are thought to be more durable than major resistance genes (McDonald and Linde, 2002; McDonald and Mundt, 2016). However, sources of partial resistance are much more difficult to identify, as subtle differences in disease severity must be accurately quantified under field conditions, ideally over time.

Automated image analysis has shown great potential to accurately quantify STB resistance and characterize different components of resistance in genetically diverse breeding material (Karisto et al., 2018; Stewart et al., 2016). However, such measurements are more labor-intensive than visual scorings and do not provide the necessary throughput to routinely screen large breeding trials over time. Some recent work has investigated the potential of reflectance-based techniques to detect and quantify STB non-destructively at the leaf and canopy level (Odilbekov et al., 2018; Yu et al., 2018). At the canopy level, the above-mentioned challenges are particularly pronounced in the case of STB, because epidemics frequently reach damaging levels and affect crop performance most during the grain filling phase (Bancal et al., 2007). Consequently, detection and quantification of STB must be achieved in fully developed canopies with a complex architecture and a clear delineation of STB and physiological senescence is essential for efficient selection.

Recently, efforts have been made to increase the specificity of reflectance-based methods. For example, new spectral vegetation indices (SVIs) with improved specificity to diseases have been developed by several authors for various patho-systems (*e.g.* Ashourloo et al., 2014; Mahlein et al., 2013). Other work has demonstrated that SVI combinations may allow to differentiate between diseases (Mahlein et al., 2010) and to delineate disease symptoms and nitrogen deficiency in wheat (Devadas et al., 2015). Yu et al. (2018) investigated the potential of different spectral features to robustly estimate STB severity at the canopy level in a large population of genetically diverse wheat genotypes. Other work has demonstrated that the sequence of temporal changes in hyperspectral reflectance signatures at the leaf level may be disease-specific, allowing to differentiate between sources of biological stress (Mahlein et al., 2012, 2010; Wahabzada et al., 2015).

Here, we aimed to achieve robust reflectance-based detection and quantification of STB under field conditions by exploiting changes in hyperspectral canopy reflectance over time. The basic rationale of the proposed approach is that pathogenesis consists in a specific and fixed sequence of events producing a constant outcome (i.e. disease symptoms). Accordingly, these events and outcomes should result in a specific and fixed sequence of changes in canopy spectral reflectance over time, irrespective of the genotype or environment under study. It seems highly likely that relying exclusively on this type of information increases the robustness of resulting estimations.

To test the feasibility of this approach, we engineered spectral-temporal features based on hyperspectral time series measurements. These features are designed to capture relevant changes in reflectance over time whilst minimizing the effect of the known confounding factors discussed above. We put forward the following hypotheses: (H_1_) Confounding effects of contrasting morphology, canopy cover and canopy 3-D structure are strongly reduced, if only relative changes in reflectance over time at the level of individual plots are analyzed. (H_2_) Confounding effects of phenology can be reduced by using combinations of STB-sensitive and STB-insensitive spectral features. Specifically, we hypothesize that several plant traits are relatively unaffected in their temporal dynamics by STB. Thus, related spectral features can be used as a baseline of changes in spectral reflectance over time, arising primarily from advancing crop phenology. This baseline can then be used to correct temporal patterns observed in STB-sensitive features for variation in phenology. Finally, we hypothesized (H_3_) that the sequence and the dynamics of STB-sensitive features is to a certain extent specific to this disease and not related to other biotic or abiotic stresses.

Thus, the objective of this study was i) to evaluate the potential of time-resolved hyperspectral reflectance measurements to detect and quantify STB infections, ii) to delineate STB and physiological senescence and iii) to estimate the robustness of the proposed method and hence its potential for breeding applications.

## 2 Materials and Methods

### 2.1 Plant Materials, Experimental Design, Phenology and meteorological data

A field experiment was carried out in the field phenotyping platform (FIP, Kirchgessner et al., 2017) at the ETH Research Station for Plant Sciences Lindau-Eschikon, Switzerland (47.449N, 8.682E, 520 m a.s.l.; soil type: eutric cambisol) in the wheat growing season of 2017-2018. A subset of 18 bread wheat (*Triticum aestivum*) genotypes was selected from the GABI wheat panel (Kollers et al., 2013; complemented with Swiss cultivars) for contrasting levels of resistance to STB and for contrasting stay-green properties based on previous experiments at the same location (Anderegg et al., 2019, *submitted to this issue*; Karisto et al., 2018). The set comprised morphologically diverse genotypes (e.g. awned and unawned) and there were obvious differences in canopy structural parameters (e.g. flag leaf angle) among the selected genotypes. The study was conducted as a two-factorial experiment in a split-plot design with the presence/absence of artificial pathogen inoculation as a whole-plot factor and genotype as a split-plot factor.

Artificial inoculation with *Z. tritici* spore suspension was done on May 21, 2018. *Z. tritici* strains ST99CH_1A5, ST99CH_1E4, ST99CH_3D1, ST99CH_3D7 were used (Zhan et al., 2002; see also http://www.septoria-tritici-blotch.net/isolate-collections.html). Spores were grown for six days in 200ml of YSB liquid media (10g yeast extract and 10g sucrose in 1l water) in several flasks for each strain. The spore suspension was filtered and pooled together for each strain. Spore concentration was adjusted and spores suspensions of each strain were mixed to achieve 150ml of inoculum for each plot containing in total 10^6^ spores/ml (2.5 x 10^5^ sp/ml of each strain). Inoculum was sprayed in the evening into wet canopy of each plot.

There were two replications for the whole-plot factor. On the same site, the entire GABI panel was also grown in two replicates (two spatially separated lots). One replication of the inoculated plots each was located in a row adjacent to a lot of the GABI panel, separated by one row sown with the resistant cultivar CH NARA (DSP, Delley, Switzerland). One replication of the non-inoculated control plots each was spatially randomized within a lot of the GABI panel. The experiments were sown with a sowing density of 400 plants m^−2^ on Oct 18, 2017. The plots sown with the GABI panel (and thus control plots within it) were treated with fungicides on three occasions: (i) Input (Bayer) was applied with a dose of 1.25 liter/ha on 23 April, 2018 (BBCH 31), (ii) Aviator Xpro (Bayer) was applied with a dose of 1.25 liter/ha on 14 May, 2018 (BBCH 51), and (iii) Osiris (BASF) was applied with a dose of 2.5 liter/ha on 28 May, 2018 (BBCH 65). The inoculated control plots did not receive fungicide treatment. Temperature data was obtained from an on-site weather station. Rainfall data was obtained from a nearby weather station of the federal Swiss meteorological network Agrometeo (www.agrometeo.ch) located at ca. 250 m distance to the field trial. The temperature data was used to calculate growing degree-days (GDD) following

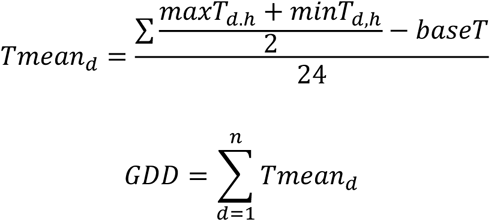

where *Tmean_d_* is the mean temperature for day d after sowing, *maxT_d,h_* and *minT_d,h_* are hourly maximum and minimum temperatures for day d after sowing and *baseT* is the base temperature, set to 0°C. Heading date was recorded when 50% of the spikes were fully emerged from the flag leaf sheath (BBCH 59, Lancashire et al., 1991). BBCH scores within the main growth stages were linearly interpolated between assessment dates. Stay-green was assessed visually as described previously (Anderegg et al., 2019, submitted), separately for the flag leaf and the whole canopy, following guidelines provided by Pask et al. (2012). Flag leaf stay-green was scored based on the portion of green leaf area on a scale from 0 (0% green leaf area) to 10 (100% green leaf area). An integer mean value was estimated for plants located in a central region of about 0.5 m × 0.5 m of each plot. Canopy stay-green was scored on the same scale by estimating the overall greenness of the plot when inspected at a view angle of approximately 45° considering the entire plot area. All scorings were done by the same person in 2-3 day intervals. An overview of measurements, scorings and samplings is given in Table 1. Growth stages were recorded until physiological maturity following the BBCH scale (Lancashire et al., 1991).

**Table 1.**
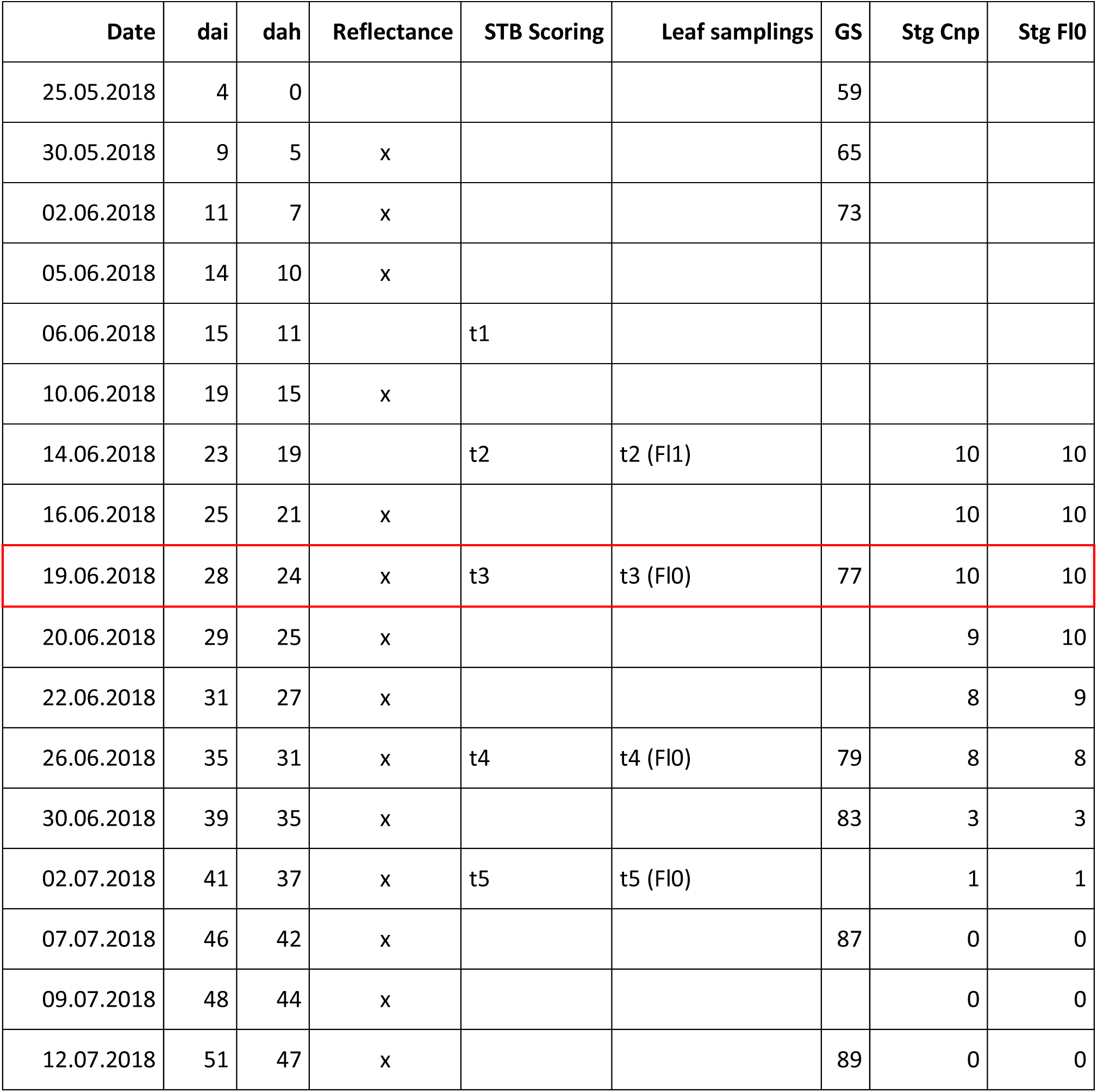
Overview of wheat phenology, canopy reflectance measurements, visual scorings and samplings. Visual incidence scorings of septoria tritici blotch (STB), Leaf samplings (Fl1 = Second leaf, Fl0 = Flag leaf), average growth stage (GS), canopy stay-green (Stg Cnp) and Flag leaf stay-green (Stg Fl0) are reported. Days after inoculation (dai) and days after heading (dah) are indicated for each date.

| Date | dai | dah | Reflectance | STB Scoring | Leaf samplings | GS | Stg Cnp | Stg FI0 |
| --- | --- | --- | --- | --- | --- | --- | --- | --- |
| 25.05.2018 | 4 | 0 |  |  |  | 59 |  |  |
| 30.05.2018 | 9 | 5 | x |  |  | 65 |  |  |
| 02.06.2018 | 11 | 7 | x |  |  | 73 |  |  |
| 05.06.2018 | 14 | 10 | x |  |  |  |  |  |
| 06.06.2018 | 15 | 11 |  | t1 |  |  |  |  |
| 10.06.2018 | 19 | 15 | x |  |  |  |  |  |
| 14.06.2018 | 23 | 19 |  | t2 | t2 (FI1) |  | 10 | 10 |
| 16.06.2018 | 25 | 21 | x |  |  |  | 10 | 10 |
| 19.06.2018 | 28 | 24 | x | t3 | t3 (FI0) | 77 | 10 | 10 |
| 20.06.2018 | 29 | 25 | x |  |  |  | 9 | 10 |
| 22.06.2018 | 31 | 27 | x |  |  |  | 8 | 9 |
| 26.06.2018 | 35 | 31 | x | t4 | t4 (FI0) | 79 | 8 | 8 |
| 30.06.2018 | 39 | 35 | x |  |  | 83 | 3 | 3 |
| 02.07.2018 | 41 | 37 | x | t5 | t5 (FI0) |  | 1 | 1 |
| 07.07.2018 | 46 | 42 | x |  |  | 87 | 0 | 0 |
| 09.07.2018 | 48 | 44 | x |  |  |  | 0 | 0 |
| 12.07.2018 | 51 | 47 | x |  |  | 89 | 0 | 0 |

### 2.2 Hyperspectral reflectance measurements

Canopy hyperspectral reflectance in the optical domain from 350 to 2500 nm was measured using a passive non-imaging spectroradiometer (ASD FieldSpec^®^ 4 spectroradiometer, ASD Inc., USA) equipped with an optic fiber with a field of view of 25°. Five spectra were recorded as the average of 15 – 25 separate spectral records while moving the fiber optic once along the diagonal of each plot at a height of approximately 0.4 m above the canopy. A Spectralon^®^ white reference panel was used for calibration before measuring canopy reflectance. The calibration was repeated after measuring one-half of a replicate (i.e. after 9 plots, approximately every 3-5 min). Measurements were carried out on 14 dates between heading and physiological maturity (i.e. between May 30 and July 12, 2018) resulting in an average of one measurement every three days. The maximum distance between two consecutive measurement dates was six days. In parallel, one lot of the GABI panel was measured on 13 dates. Here, the sensor calibration was repeated approximately every 10 min after completion of measurements on two rows.

### 2.3 STB disease assessment

The amount of STB in each plot was assessed on five dates (t1 – t5) between 16 days after inoculation (dai, June 6, 2018) and 42 dai (July 2, 2018). STB was quantified by combined assessments of disease incidence (i.e. the proportion of leaves showing visible symptoms of STB) and conditional disease severity (i.e. the amount of disease on symptomatic plants). Disease incidence was assessed visually for 30 plants per plot by inspecting the leaves of one tiller per plant. Incidence scorings were obtained per leaf layer. Conditional disease severity was then measured using automated analysis of scanned leaves exhibiting obvious disease symptoms. To this end, eight infected leaves were sampled per plot, transported to the laboratory and imaged on flatbed scanners following the method described by Stewart et al. (2016) and Karisto et al. (2018). However, to avoid interfering excessively with the development of the disease epidemic, leaf samples from inoculated plots were taken only if disease incidence was at least 1/3 (i.e. if at least 10 out of 30 examined leaves exhibited symptoms of STB infection). Thus, no leaf samples were taken at t1, while at t2, second leaves from the top were sampled from a subset of plots. Starting at t3, all plots were sampled at the flag leaf layer. In contrast, from non-inoculated control plots, eight leaves were sampled without reference to their disease status due to very low disease incidence. Automated image analysis was then used to extract the percent of leaf area covered by lesions (PLACL) from the generated leaf scans using thresholds in the HSV color space and functions of the python API of openCV V3.0.0 (https://opencv.org/). The procedure was optimized to minimize the effect of insect damage, powdery mildew infections and physiological senescence, particularly leaf-tip necrosis, on the derivation of PLACL. PLACL was extracted only from t2 and t3 scans, as leaves increasingly displayed physiological senescence at later time-points. The developed python script with detailed annotations can be retrieved from github (https://github.com/and-jonas/stb_placl). Finally, overall disease severity was calculated by multiplying disease incidence with conditional disease severity for inoculated plots, whereas it was directly extracted from the leaf scans for control plots.

### 2.4 Data analysis

All data analyses were done in the R environment for statistical computing (R version 3.5.2; R Core Team, 2018). Raw spectra were smoothed using the Savitzky-Golay smoothing filter (Savitzky and Golay, 1964) with a window size of 11 spectral bands and a third order polynomial, using the r-package ‘*prospectr*’ V0.1.3. (Ramirez-Lopez and Stevens, 2014). Spectral regions comprising the wavelengths from 1350 nm to 1475 nm, from 1781 nm to 1990 nm and from 2400 nm to 2500 nm were removed because of the very low signal-to-noise ratio resulting from high atmospheric absorption. Spectra were averaged for each experimental plot. Pre-processed spectra, consisting of reflectance values at 1709 wavelengths, were then used for time-point specific analysis as well as for time-integrated analyses, as described in the next sections. For ease of notation, the reflectance at a specific wavelength will be abbreviated by R followed by the wavelength in nm (e.g. R750).

#### 2.4.1 Benchmark time-point specific analysis

The relationship between spectral reflectance and STB was studied on a diverse panel of wheat genotypes by Yu et al. (2018), but the analysis was limited to single time-points. We performed a comparable analysis for each measurement time-point as a benchmark and to estimate model transferability across time. Yu et al. (2018) reported improved prediction of STB severity metrics and classification accuracy when using the full spectrum rather than single SVIs. Therefore, our analysis focuses on these approaches. We tested two parametrically structured linear models (Partial Least Squares (PLS) regression and ridge regression) and two tree-based ensemble models (random forest regression and cubist regression) for their capability to predict STB severity metrics. For classification, Partial Least Squares Discriminant Analysis (PLSDA) was used (*for details on these methods we refer to* Kuhn and Johnson, 2013 *and citations therein*). Prior to modelling, spectral resolution was reduced to 6 nm by binning (i.e. by computing average values for six adjacent wavelengths). Following a standard procedure (Kuhn and Johnson, 2013), model hyperparameters were tuned using 10-times repeated 10-fold cross-validation, with the root mean square error of predictions (RMSE) and overall classification accuracy as performance metrics for regression and classification, respectively. The overall accuracy reflects the agreement between the predicted and the observed classes. This agreement can also be expressed in terms of sensitivity and specificity of the model, with

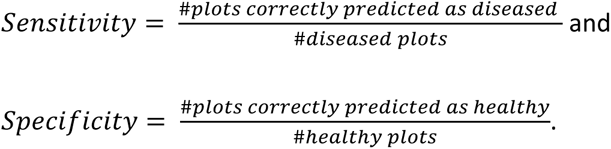

The simplest model with a performance within one standard error of the absolute best model was chosen as the final model. Variable importance for the projection (VIP) was computed for PLSDA models to estimate the importance of individual wavebands to predict the class (i.e. ‘healthy’ or ‘diseased’). In the regression setting, two different training datasets were used for model fitting: the full dataset, including all control plots, and a dataset consisting of the inoculated plots only. When all control plots were used for model fitting, the RMSE and *R*^2^ of the resulting models was adjusted by removing the predicted and observed values for the control plots again, in order to avoid overly optimistic performance estimates resulting from a good prediction of disease severity in control plots. The R-packages ‘*caret*’ V6.0.80 (Kuhn, 2008), ‘*mixOmic*s’ V6.3.2 (Rohart et al., 2017), ‘*pls*’ V2.7.0 (Mevik et al., 2018), ‘*Cubist*’ V0.2.2 (Kuhn et al., 2018), ‘*ranger*’ V0.10.1 (Wright and Ziegler, 2017) and ‘*elasticnet*’ V1.1.1 (Hastie, 2018) were used for the analysis.

#### 2.4.2 Time-integrated analysis

Summarizing H_1_-H_3_, we hypothesized that the analysis of temporal dynamics in hyperspectral reflectance signatures may facilitate a robust detection and quantification of STB across diverse wheat genotypes under field conditions. To evaluate these hypotheses, we condensed the hyperspectral time series into time series of SVIs, similar to the procedure described previously (Anderegg et al., 2019, *submitted*). Thereby, we obtained a comprehensive summary representation of the hyperspectral dataset collected over time, interpretable in terms of plant physiology and canopy characteristics. The smoothness of SVI values over time was evaluated graphically and only SVIs showing a clear and interpretable temporal trend were maintained for further analyses. Values of the selected SVIs were scaled to range from 0 to 10, representing the minimum and maximum values recorded during the assessment period for the corresponding experimental plot, respectively. To simplify subsequent steps in the analysis, the scale for SVIs with increasing values over time was inverted. Measurement dates were converted to thermal time after heading by subtracting the plot-specific accumulated thermal time at heading from the accumulated thermal time at each measurement date. The scaled SVI values were then fitted against thermal time after heading for each experimental plot and SVI using a Gompertz model with asymptotes constrained to 0 and 10 (eq. 1).

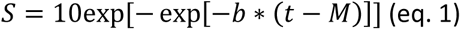

where *S* represents the scaled SVI value, *t* is the accumulated thermal time after heading, *b* is the rate of change at time *M* and *M* is the accumulated thermal time after heading when the rate is at its maximum (Gooding et al., 2000). Eq. (1) was fitted using the R package ‘*nls.multstart*’ V1.0.0 (Padfield and Matheson, 2018). Two types of dynamics parameters for each experimental plot and SVI were extracted from the resulting model fits: (1) ‘key time-points’, which are specific points in time when a certain criterion (e.g. a threshold) is met; and (2) ‘change parameters’, which represent the rate or duration of a process (Figure 1). We extracted two key time-points: the *M* parameter of the Gompertz model, and the time when fitted values decreased to 8.5 (t_85_). As change parameters we used the rate parameter *b* of the Gompertz model, and the duration between t_85_ and M (dur). While the b and M parameters of the Gompertz model fully describe the fitted curve, the t_85_ and dur parameters are affected by both Gompertz model parameters, thus representing a mix of both. The threshold was set to 8.5 because i) this level efficiently captured observed variation during the late stay-green phase (Figure 2), ii) it was little affected by somewhat unstable values during the stay-green phase observed for some SVIs, and iii) for some SVIs, the initial highest values were not optimally represented by the Gompertz model.

**Figure 1.**
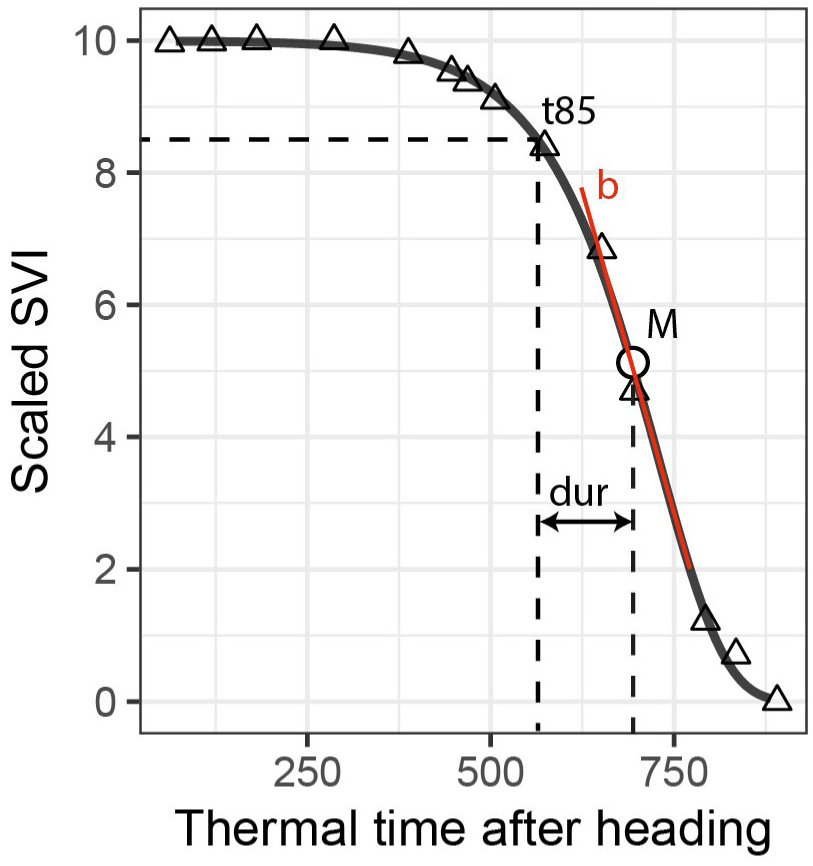
Extraction of dynamics parameters for one spectral vegetation index (SVI; here scaled values of the Plant Senescence Reflectance Index) and one experimental plot (here a non-inoculated control plot). The t<u>_85_</u> parameter is the time point when fitted scaled SVI values decrease to 8.5; M is a parameter of the Gompertz model, representing the time point when the rate of decrease is at its maximum; the dur parameter represents the duration in thermal time between t_85_ and M; b is a parameter of the Gompertz model, representing the maximum rate of decrease. M and t_85_ are labelled ‘key points’, dur and b are labelled ‘change parameters’.

**Figure 2.**
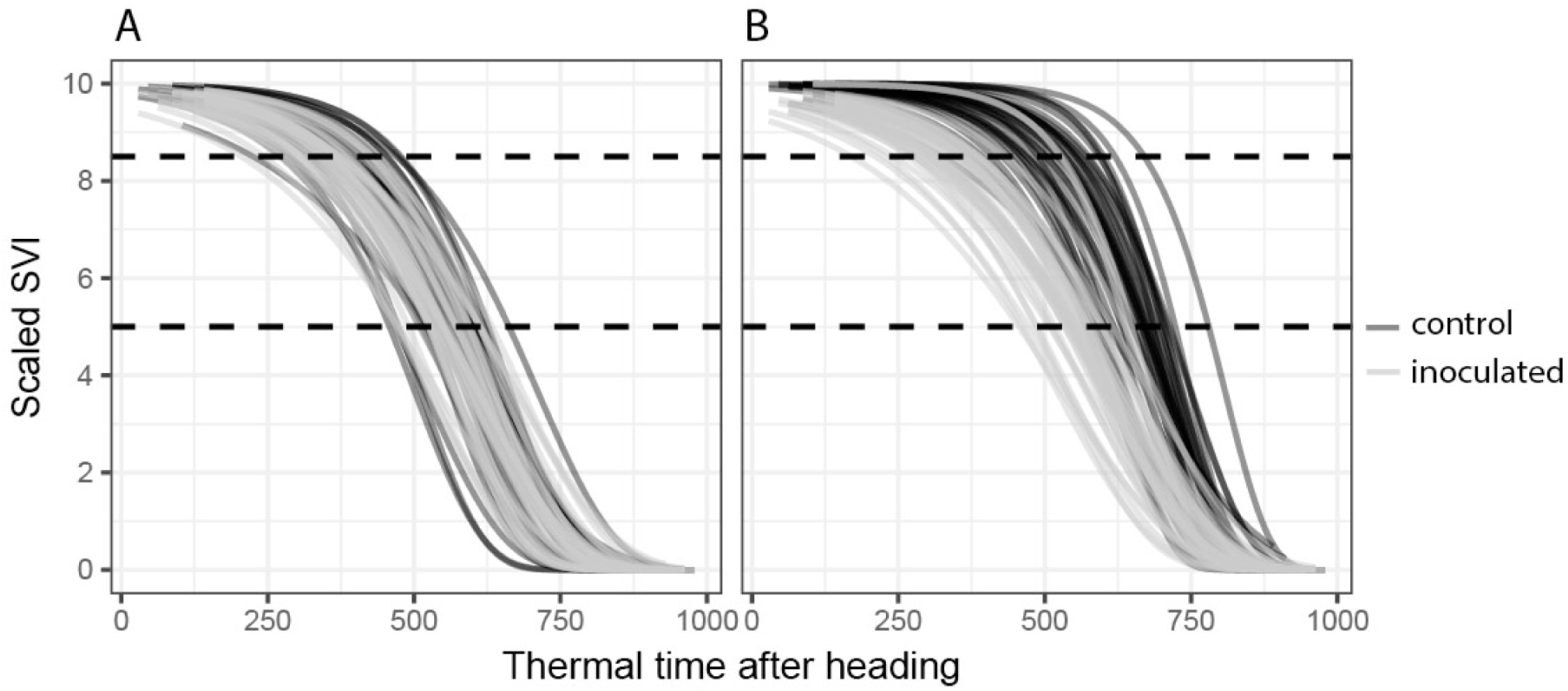
Temporal dynamics of spectral vegetation indices (SVIs). Gompertz model fits for all 72 experimental plots are shown. **(A)** A disease-insensitive SVI (here the Flowering Intensity Index, FII) displays the same temporal patterns for both treatments (inoculated and control). **(B)** A disease-sensitive SVI (here the Modified Chlorophyll Absorption Ratio Index, MCARI2) displays contrasting temporal patterns for control and inoculated plots.

Next, SVIs least affected in their temporal dynamics by the presence or absence of STB infections were selected separately for each dynamics parameter as follows: for the key time-points (t_85_, M) by selecting SVIs with the smallest average difference between the parameter values of the inoculated and non-inoculated control plots; for change parameters (b, dur) by selecting SVIs with the smallest average deviance from 1 of the ratio of the change parameter values. For each dynamics parameter, a subset of eight SVIs with the smallest difference or ratio was selected. All other SVIs were considered to be significantly affected by STB infection. Figure 2 shows an example of an STB-sensitive and an STB-insensitive SVI.

We then performed unsupervised subset selection (i.e. without considering the response) on both sets of SVIs (i.e. the STB-sensitive and STB-insensitive SVIs) with the aim of removing redundant SVIs. For each dynamics parameter (t_85_, M, dur, b), pairwise Pearson correlation coefficients between the parameter values derived from all used SVIs were computed. For change parameters, the maximum linear correlation allowed was set to r = 0.9, whereas for the key time-points it was set to r = 0.95, as these were generally highly collinear. In cases where pair-wise correlations were higher than these threshold values, only one of the two SVIs was retained, preferring narrow-band SVIs over broad-band SVIs, SVIs with a specific physiological interpretation and SVIs developed specifically for use in wheat or barley canopies over more generic SVIs. Additionally, the goodness of the Gompertz model fit was evaluated qualitatively (i.e. graphically) and used as an additional selection criterion. The parameters were then combined by calculating differences between the key time-points derived from selected STB-sensitive and STB-insensitive SVIs and the ratios of the change parameters derived from STB-sensitive and STB-insensitive SVIs (Figure 3). These differences and ratios were calculated for all possible pairs of STB-sensitive and STB-insensitive SVIs and were then used as features for (1) the classification of plots into non-inoculated healthy control plots and inoculated, diseased plots and (2) the prediction of STB severity in each plot. Models and model fitting procedures were identical to the time-point specific analysis.

**Figure 3.**
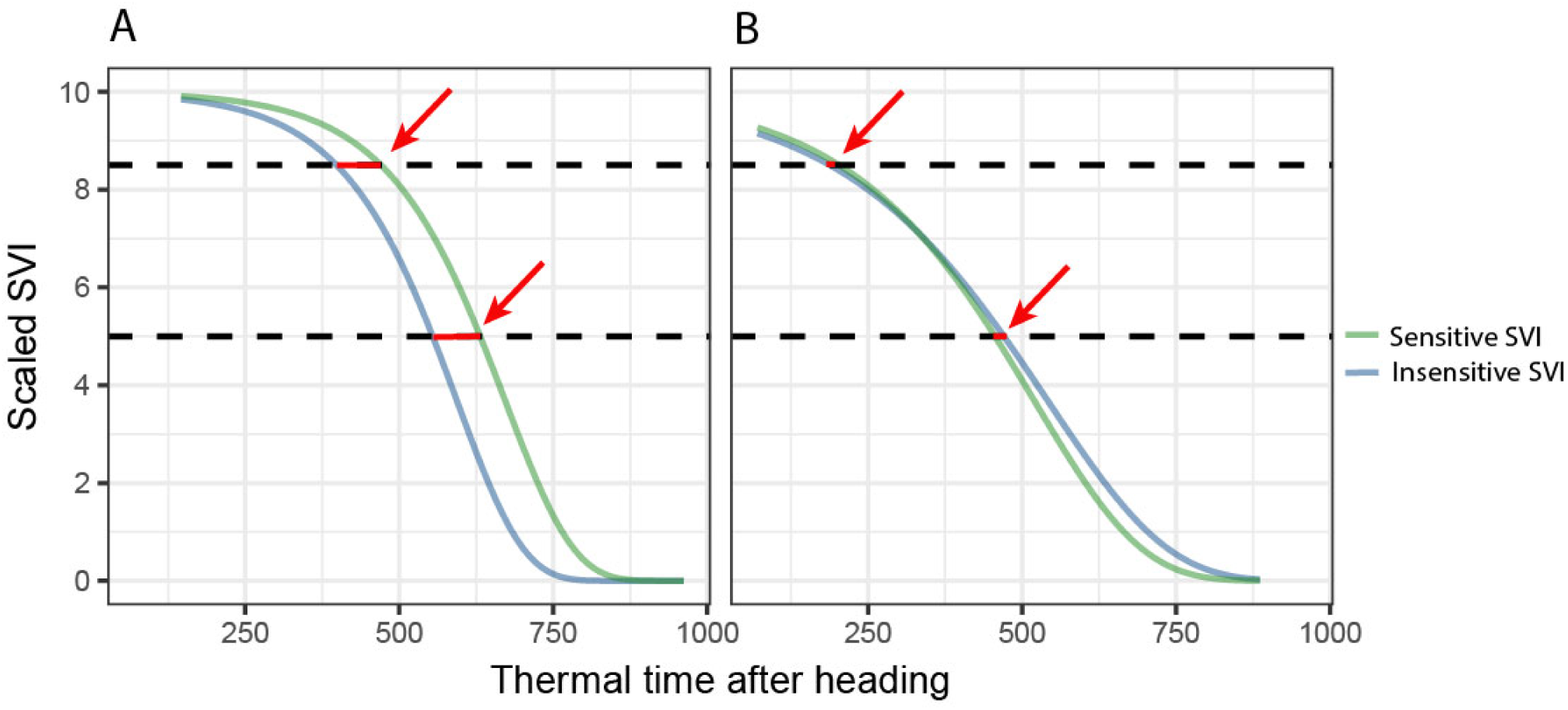
Derivation of the final key time points based predictors for disease classification and quantification. Key time points extracted from disease-sensitive and disease-insensitive spectral vegetation indices (SVI) are combined to isolate the effect of the disease from other effects (e.g. contrasting stay-green) by calculating the differences (highlighted by red arrows). **(A)** Control, ‘healthy’ plot, **(B)** Inoculated, ‘diseased’ plot. For change parameters, the ratio, rather than the difference, was calculated.

#### 2.4.3 Selection of spectral-temporal features

While tree-based models are considered naturally resistant to non-informative predictors, and some perform feature selection intrinsically, the presence of highly correlated predictors makes the interpretation of resulting variable importance measures challenging (Strobl et al., 2007). Hence, we performed supervised feature selection by recursive feature elimination with cubist and random forest regression as base-learners, using a nested cross-validation approach. The dataset was resampled 30 times with an 80:20 split using stratified sampling. Samples were binned into eight classes based on percentiles of STB severity to ensure balanced evaluation datasets. Thus, for each resample, feature elimination was carried out on 80% of the data, and model performance was evaluated on the remaining 20% in 28 decreasing steps. Eliminated features were assigned a rank corresponding to the iteration after which they were excluded (i.e. those eliminated first had rank 28, whereas the feature retained as the last had rank 1). In each iteration, the base-learner hyperparameters were tuned using 10-fold cross-validation (*see* Ambroise and McLachlan, 2002; Granitto et al., 2006; Guyon et al., 2002 *for a detailed discussion of the methodology*). The analysis was implemented using a custom-developed R-Script (https://github.com/and-jonas/rfe).

#### 2.4.4 Independent model validation

The performance and robustness of the developed models was further evaluated using data of 360 wheat plots obtained from one replication of the GABI panel. Low to intermediate disease incidence and very low conditional disease severity as well as late appearance of symptoms in all 36 control plots spatially randomized within the two replications of the panel suggested that STB disease should not have reached damaging levels in the vast majority of these plots. Therefore, these plots were considered as essentially disease-free. For all of these plots, spectral-temporal features were extracted from the 13 measurement time-points as described previously and were then used to generate a class label and class probabilities from the classification models as well as a prediction of disease severity from the regression models. To distinguish the performance measures obtained for held-out samples of the main experiment (i.e. the cross-validated training performance) from those obtained for the independent plots, these are referred to as the internal accuracy (acc_int_) and the generalized accuracy (acc_gen_), respectively. It should be noted that accuracy represents only model specificity in this case, as no independent plots with significant levels of STB were available. In a final validation step, the spatial distribution of class labels and severity predictions were examined by creating plots of the experimental design. Thus, we aimed to test the robustness of the models to heterogeneous field conditions. Field heterogeneity may affect plant physiology and thus hyperspectral reflectance over time (e.g. through the development of local drought stress). The presence or absence of spatial patterns in model predictions can therefore be interpreted as an additional measure of model robustness.

## 3 Results

### 3.1 Development of STB disease

Towards the end of the vegetation period, all inoculated plots had substantial levels of STB. In contrast, non-inoculated control plots were essentially disease-free until late in the vegetation period. Thus, artificial inoculations were effective in all plots and the dataset was suitable for testing the feasibility of the classification of plots into healthy and diseased canopies based on reflectance spectra or spectral-temporal features. Furthermore, large variation in the levels of STB could be observed among the inoculated plots, probably attributable to different levels of resistance, with the largest variation observed during late stay-green (i.e. at t3, June 19, 2018, compare with Figure 4B). Thus, the dataset was also suitable for testing the feasibility of disease quantification using reflectance spectra or spectral-temporal features.

**Figure 4.**
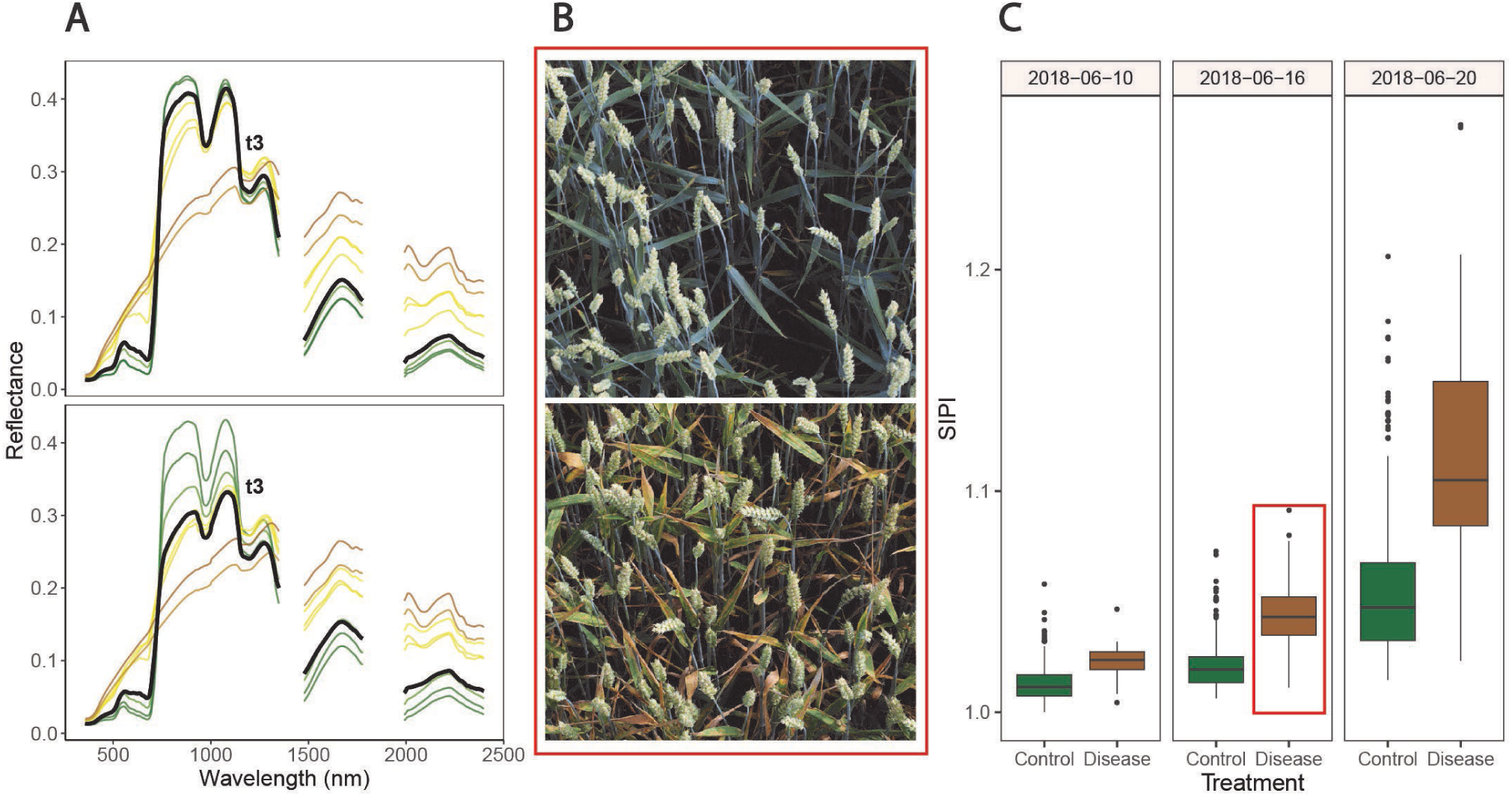
Symptoms of Septoria tritici blotch (STB) and associated spectral reflectance characteristics over time. **(A)** Date-wise averaged reflectance spectra of healthy canopies (upper panel) and of inoculated, diseased canopies (lower panel). Colors approximate the average color of the vegetation on the corresponding measurement date (estimated based on average visual canopy senescence scorings). The thick black spectra mark the average reflectance spectra measured at t3 (i.e. June 19, 2018). **(B)** Images of two inoculated plots, taken on June 16, 2018. The genotype in the upper panel was highly resistant to STB, developing visible symptoms only later, whereas the genotype in the lower panel was highly susceptible and displays severe symptoms of STB. Images were captured using the Field Phenotyping Platform (FIP, Kirchgessner et al., 2017). **(C)** Values of the Structure Insensitive Pigment Index (SIPI) for both treatments on three measurement dates. The plots shown in panel B are contained in this boxplot (indicated by the red box). No obvious signs of apical senescence were visible by June 20, 2018 in any of the healthy control plots, but senescence started shortly after.

High levels of STB in inoculated plots were the result of both high incidence and high conditional severity, particularly at t4 and t5 (Figure 5A, Table 2). Visual assessments of scanned leaves suggested a high conditional severity in all inoculated plots at these late stages. In contrast, the non-inoculated control plots displayed very low levels of STB even at late stages. STB incidence increased in some plots at t4 and t5, but visual assessments of the sampled leaves demonstrated very low conditional disease severity even at t5. Since the subset of genotypes used for the experiment also included some highly susceptible genotypes, this suggests that natural infections did not cause agronomically significant levels of STB in this experiment. This was likely the result of fungicide applications and the very low rainfall in the period from May to July. Rainfall in this period totaled 178 mm, which represents 52% of the average of 343 mm in the reference period 1981–2010 (MeteoSwiss, 2019).

**Figure 5.**
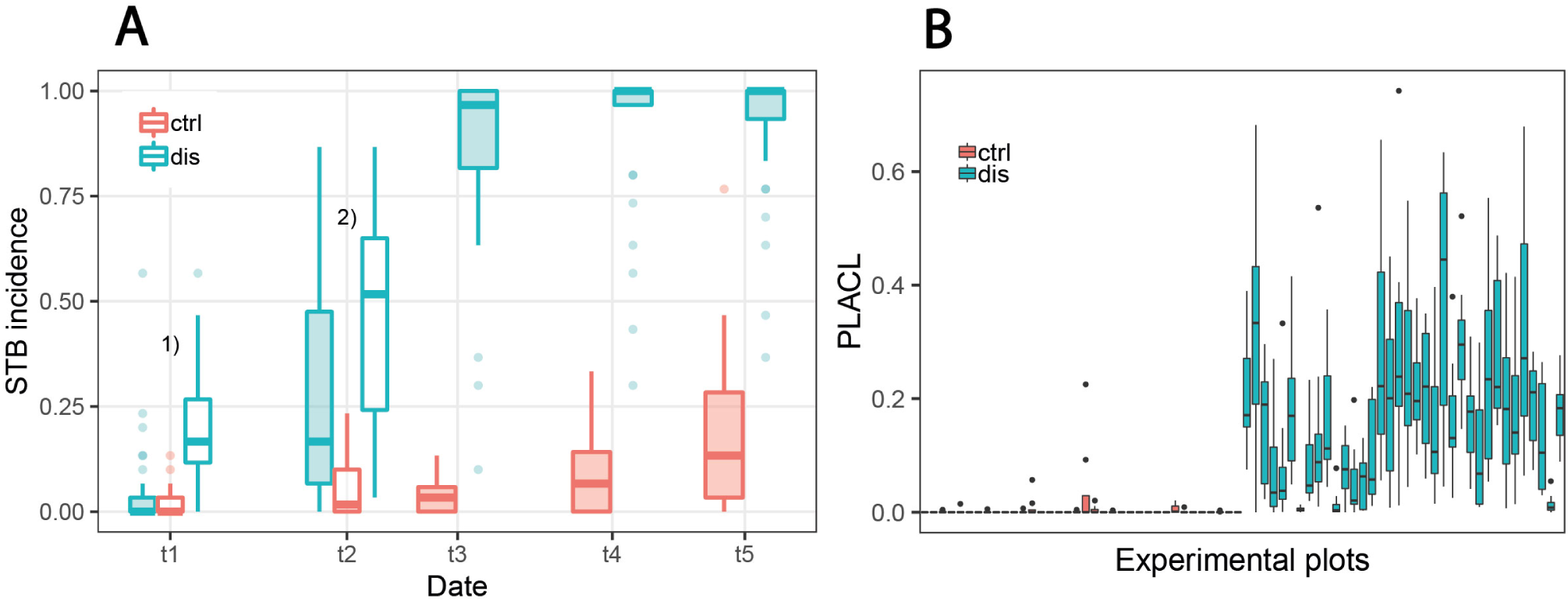
Development of septoria tritici blotch (STB) disease. **(A)** STB incidence at five different assessment dates for non-inoculated ‘healthy’ control plots (ctrl) and for artificially inoculated ‘diseased’ plots (dis). In diseased plots, STB incidence on flag leaves was assessed at all time points, whereas for the control plots, it was assessed only from t3 onwards. Filled boxes represent STB incidence on flag leaves; open boxes represent STB incidence on lower leaf layers. ^1)^Open boxes represent STB incidence on third leaves from the top, ^2)^open boxes represent STB incidence on second leaves from the top. **(B)** Conditional disease severity measured as percent leaf area covered by lesions (PLACL) at t3 (June 19, 2018) for eight flag leaves per plot for all 72 experimental plots.

**Table 2.** Summary of septoria tritici blotch (STB) assessments. STB incidence and conditional severity was assessed at the level of individual leaf layers, namely the flag leaf (Fl0), the penultimate leaf (Fl1) and the ante-penultimate leaf (Fl2). Conditional severity was measured as percent leaf area covered by lesions (PLACL). Severity was calculated as the product of STB incidence and conditional severity. Values are reported separately for non-inoculated control plots and inoculated plots, separated by a slash. Mean values across all plots are reported, with minima and maxima in brackets. Disease assessments were carried out on five dates (t1 – t5) covering the growth phases of 15 days after inoculation (dai) to 41 dai.

|  |  | STB Incidence |  |  | PLACL |  | Severity |  |
| --- | --- | --- | --- | --- | --- | --- | --- | --- |
| Time point | dai | FI0 | FI1 | FI2 | FI0 | FI1 | FI0 | FI1 |
| t1 | 15 | - /<br>0.05 (0, 0.57) | - /<br>0.10 (0, 0.53) | 0.02 (0, 0.13) /<br>0.20 (0, 0.57) | - /<br>- | - /<br>- | - /<br>- | - /<br>- |
| t2 | 23 | - /<br>0.26 (0, 0.87) | 0.05 (0, 0.23) /<br>0.45 (0.03, 0.87) | - /<br>- | - /<br>- | - /<br>0.05 (0, 0.25) | - /<br>- | - /<br>0.03 (0, 0.20) |
| t3 | 28 | 0.03 (0, 0.13) /<br>0.87 (0.10, 1) | 0.09 (0, 0.50) /<br>0.91 (0.10, 1) | - /<br>- | - /<br>0.17 (0, 0.38) | - /<br>- | **0 (0, 0.04) /**<br>**0.16 (0, 0.38)** | - /<br>- |
| t4 | 35 | 0.09 (0, 0.33) /<br>0.92 (0.30, 1) | - /<br>- | - /<br>- | - /<br>- | - /<br>- | - /<br>- | - /<br>- |
| t5 | 41 | 0.17 (0, 0.77) /<br>0.92 (0.37, 1) | - /<br>- | - /<br>- | - /<br>- | - /<br>- | - /<br>- | - /<br>- |

STB incidence was low at t1, both in inoculated and in control plots. Symptoms were apparent at significant levels only on lower leaf layers of inoculated plots, whereas flag leaves were essentially disease-free in both treatments. At t2, there was approximately a five-fold increase in STB incidence at the flag leaf and subtending leaf layer in many inoculated plots (Figure 5A, Table 2). At t3, STB incidence on flag leaves reached very high levels in most inoculated plots, and PLACL reached an average of 17%, indicating a moderate conditional STB severity on average. Thus, the observed differences in STB severity among inoculated plots is primarily a result of differences in conditional severity, with PLACL ranging from 0% to 38% (Figure 5B). In control plots, almost no lesions were detected. There were some signs of physiological senescence on sampled flag leaves at t3, but these were mostly limited to yellowing of the entire leaves and/or leaf tip necrosis. As there was ample variation for disease severity at t3 among the inoculated plots, and physiological senescence did not significantly affect extraction of PLACL from leaf scans, this time-point was chosen as a measure of overall disease severity and used as response variable in the time-integrated analysis.

### 3.2 Effects of STB and phenology on canopy spectral reflectance

Over the assessment period, observed changes in spectral reflectance were similar for diseased and healthy canopies (Figure 4A), showing the typical pattern of senescing canopies. However, an obvious effect of STB infections consisted in an early marked decrease in reflectance in the NIR not observable in healthy canopies. This decrease preceded the appearance of physiological senescence and was observable in the pre-symptomatic phase of STB infections. Furthermore, an early increase in reflectance in the VIS, especially for wavelengths greater than 535 nm, was observed. An early increase in SWIR reflectance in diseased canopies compared to healthy canopies was also discernable. However, these differences were small compared to changes in reflectance over time.

Canopy spectral reflectance seemed to remain relatively constant throughout the stay-green phase in healthy canopies (Figure 4A, upper panel). However, the examination of SVI values over time revealed significant changes in canopy reflectance during this period (Figure 4C). Importantly, variation caused by advancing phenology (i.e. within-treatment variation in Figure 4C) was prominent with respect to STB-induced variation (i.e. between-treatment variation in Figure 4C). This is true even for the structure insensitive pigment index (SIPI), which has been proposed as a potential surrogate for crop disease under field conditions (Yu et al., 2018; Figure 4C). For several other spectral indices, initial variation as well as variation over time was even larger (data not shown).

### 3.3 Time-point specific full-spectrum analysis

#### 3.3.1 Binary classification into healthy and diseased canopies using reflectance spectra

PLSDA models correctly classified all held-out samples in most resampling iterations, resulting in classification accuracies acc_int_ ≥ 0.96 for all time-points (Figure 6). The optimal number of components used by the PLSDA models (determined via repeated CV) was between 5 and 17, depending on the time-point.

**Figure 6.**
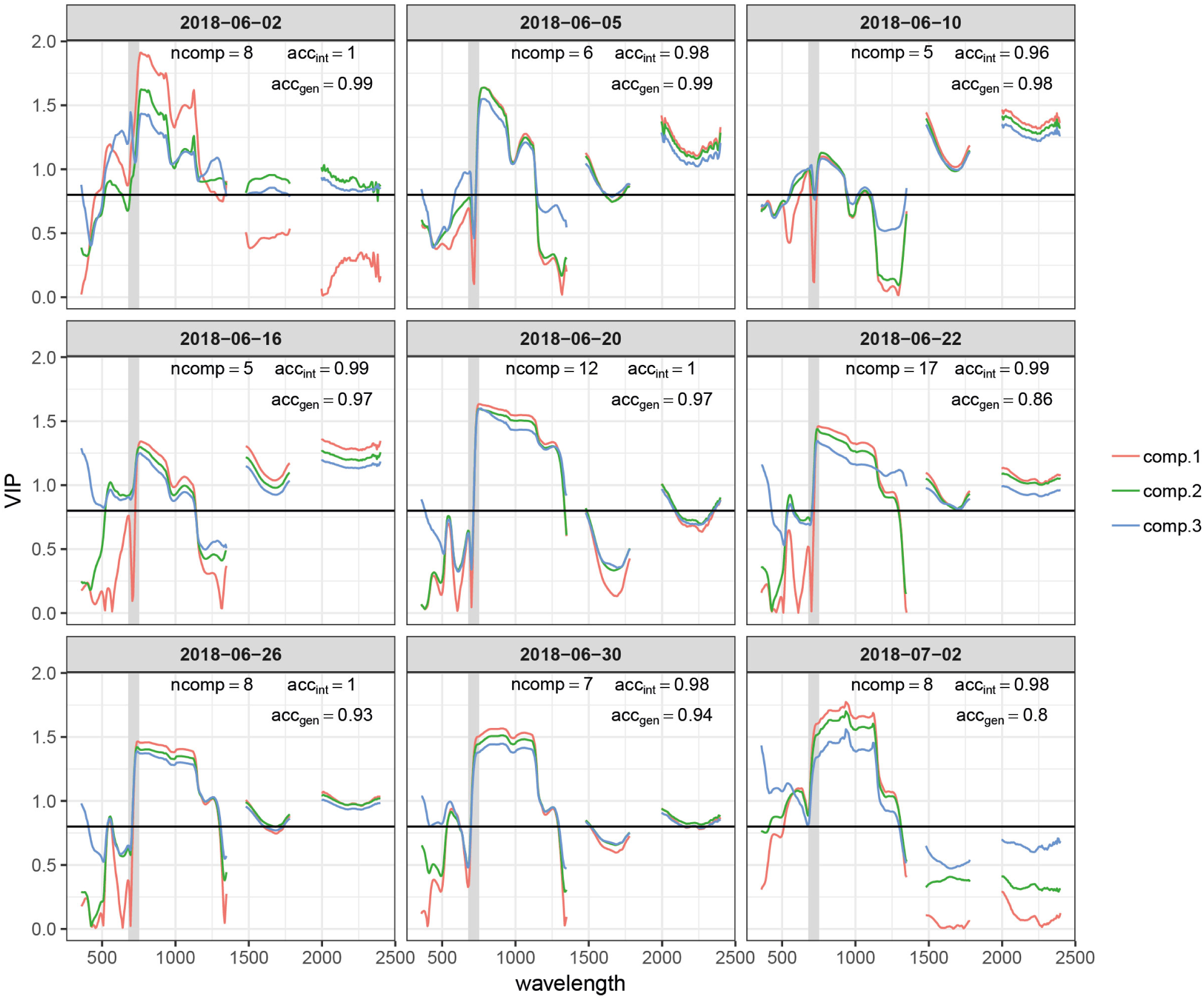
Variable importance for the projection (VIP) of the time point specific partial least squares discriminant analysis (PLSDA) models for the first three components (comp.1 – comp.3). The total number of components used by the model (ncomp), the prediction accuracy for plots included in the experiment (acc_int_) and prediction accuracy for the independent test set comprising >300 plots of an adjacent experiment measured on the same day or with a maximum delay of one day (acc_gen_) are also reported for each model. The earliest and latest time points are not represented. The grey shaded area represents the spectral range between 680 nm and 750 nm, i.e. the red edge. The horizontal black line marks a commonly used threshold value for an important contribution (VIP = 0.8).

Correct class labels were obtained for all held-out samples even for the first time-point at 9 dai, when no visual symptoms of STB were present in most experimental plots. However, prediction accuracies for the independent GABI plots were distinctly lower, particularly for models calibrated with data from early and late measurement time-points (data not shown). A satisfactory performance on independent plots was observed for models calibrated with data from the late stay-green phase (i.e. between 2018-06-10 and 2018-06-22), which correctly predicted the independent plots as disease-free in most cases (acc_gen_ ≥ 0.88 in all cases).

VIP scores quantify the importance of wavebands to predict the response, i.e. to generate the class label (‘healthy’ or ‘diseased’) or to predict STB disease severity here (Yu et al., 2018). VIP scores for the first three components showed some general patterns across time-points (Figure 6). The near-infrared region (NIR, 750-1300 nm) and the short-wave infrared region (SWIR; 1475-1781 nm and 1991-2400 nm) had a relatively high importance (Figure 6). However, the relative importance of the SWIR compared to the NIR drastically changes over time. The importance of the SWIR is comparably low during early stay-green, but its importance greatly increases and exceeds the importance of the NIR during late stay-green. Furthermore, the red-edge (RE, 680-750nm) had a low importance at the beginning, but is increasingly important at later stages, as indicated by a gradual left-shift of the peak in VIP at the NIR for later time-points. Finally, at early time-points, there is a significant contribution of wavebands in the visible range (VIS, 400-700nm). This feature is somewhat transformed over time, resulting in a narrow peak in VIP at wavelengths around 535nm at intermediate time-points. Towards later time-points, this feature broadens again.

Given the common patterns but also significant differences across time-points, we aimed to evaluate the robustness of the developed models to temporal changes in reflectance induced by advancing crop phenology, as differences in crop phenology are typically present among genotypes in breeding programs. Model performance across time is shown in Figure 7. The performance of models calibrated with data from early and late time-points quickly deteriorates. In contrast, models calibrated with data of intermediate time-points show a higher stability over time, although the performance of some models still decreases rather fast. The models created using data from 2018-06-10 and 2018-06-19 were most robust over time, and produce accurate class predictions over a period of about 10 days. It is essential to note that these performance estimates are derived from the same plots used to calibrate the models, although at different time-points. Given the lower acc_gen_ compared to the acc_int_ (see above), significantly decreased performance should be expected on entirely independent plots (different genotypes).

**Figure 7.**
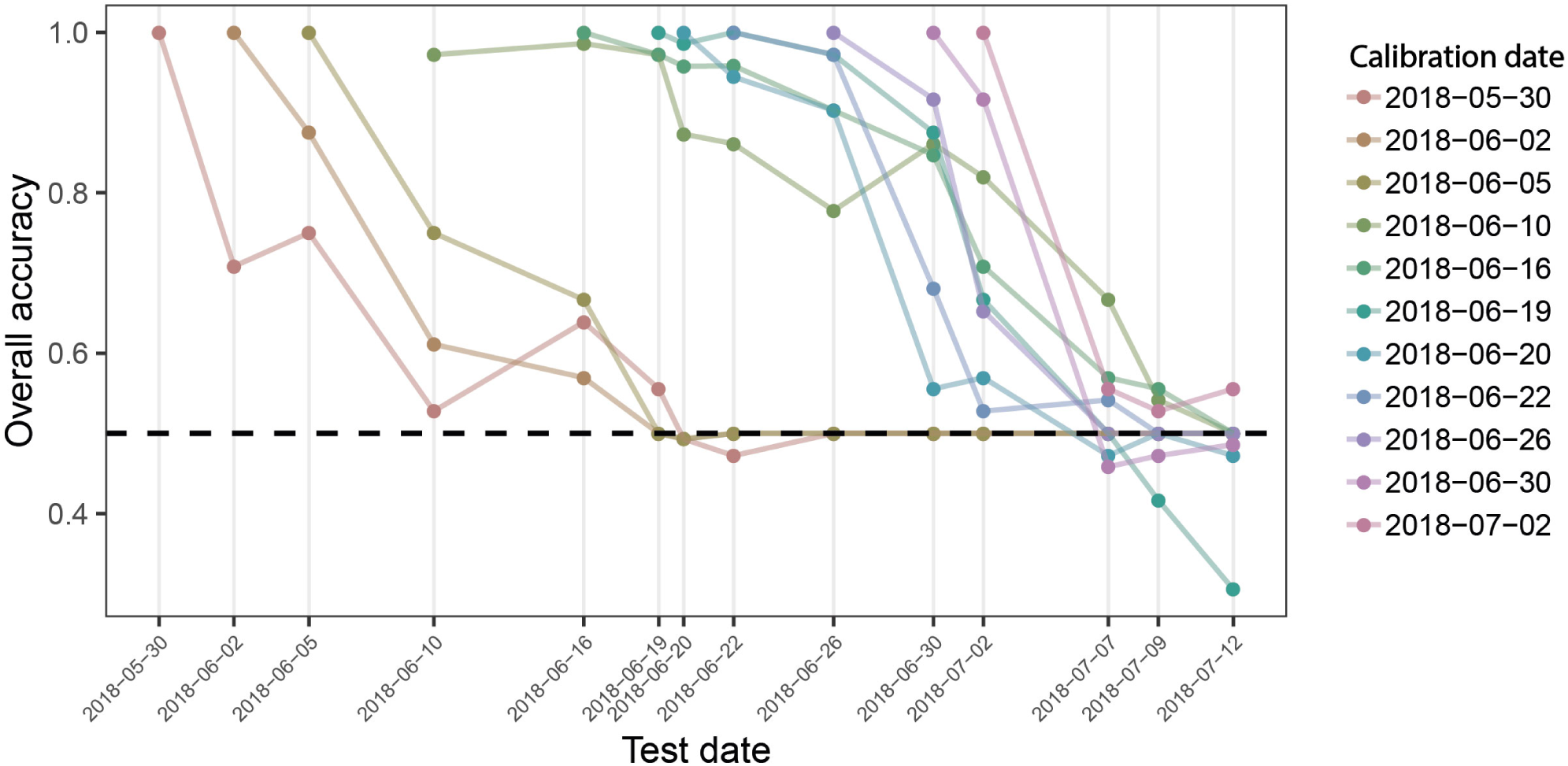
Overall prediction accuracy of PLSDA models across time. Binary classification models were calibrated for each measurement time point, using reflectance spectra collected on this date as predictors and the known class label (i.e. “diseased” or “healthy”) as response. The performance of these models was evaluated on held out samples of the same date as well as on all plots of the subsequent measurement time points as the overall accuracy of classification. Colored lines track the performance of each date-specific model across time (e.g. the left-most red line represents the performance of the model calibrated with spectra obtained on May 30, 2018, when tested on the same date, and for all subsequent measurement dates). The broken black line indicates a performance of 0.5, i.e. the performance of a random guess of the class label.

#### 3.3.2 Regression models to quantify disease severity using reflectance spectra

Cubist regression models performed best in predicting disease severity. The smallest RMSE was obtained for models trained on data from inoculated plots only (RMSE = 0.061, *R*^2^ = 0.67; Figure 8). The underlying model was simple, building on only four variables (R748, R766, R892 and R1084) in a single model tree. Model performance was slightly decreased when all available data was used for model fitting (RMSE_adj_ = 0.066, *R*^2^_adj_ = 0.55). Here, significant improvements were achieved by increasing model complexity. Validation on the largely disease-free plots of the GABI panel suggested a high specificity of the model (i.e. disease levels on healthy plots were predicted to be virtually zero for almost all plots). This was true irrespective of whether the control plots were included in the training dataset or not. PLS and ridge regression performed similarly (RMSE = 0.063, *R*^2^ = 0.64), whereas random forest regression performed comparably poorly (RMSE = 0.087, *R*^2^ = 0.46). A strong systematic bias was observed in predictions of the random forest, with low values of disease severity overestimated and high values underestimated.

**Figure 8.**
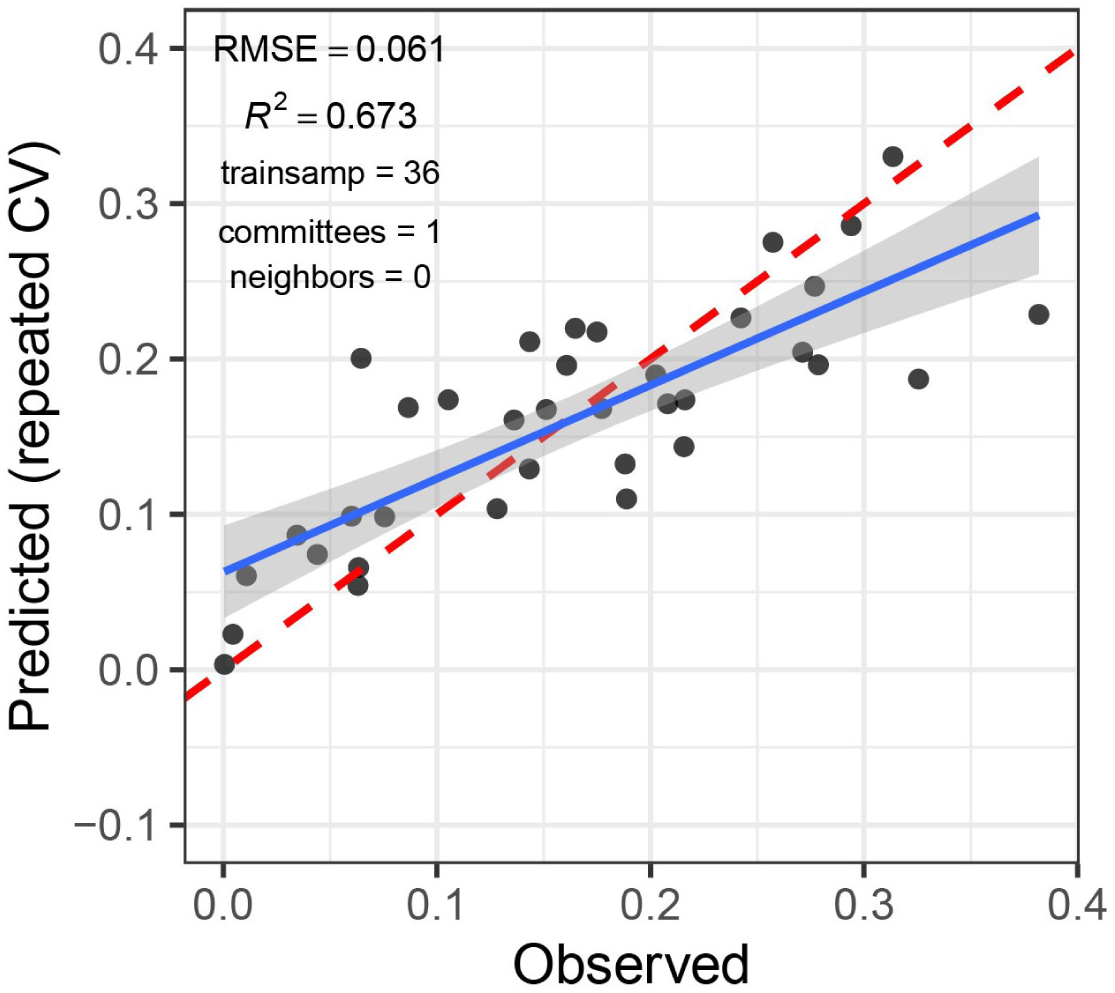
Predicted vs. observed septoria tritici blotch (STB) severity levels of 36 artificially inoculated wheat plots. STB severity was measured on flag leaves using a combination of visual incidence scorings and scans of flag leaves exhibiting disease symptoms. Predictions were obtained from a cubist regression model based on the reflectance spectrum of the canopies measured on June 19, 2018.

### 3.4 Time-integrated analysis using combinations of spectral vegetation indices

#### 3.4.1 Engineering of spectral-temporal features as new predictors

Fifty-seven of the tested SVIs were deemed amenable for analysis of their temporal dynamics in the proposed framework (i.e. they displayed a clear and interpretable temporal trend, which could be represented using a Gompertz-type model). For seven of these, the last three measurements were excluded prior to modelling their temporal dynamics, as values increased again in later stages. After subset selection, seven and 13 SVIs were retained as insensitive and sensitive SVIs, respectively, for the key time-points. For change parameters, four and twelve SVIs were retained as insensitive and sensitive SVIs, respectively. In total, 24 distinct SVIs were retained, of which ten sensitive and 14 insensitive SVIs. From their fitted dynamics, a total of 278 (i. e. 4 insensitive SVIs * 12 sensitive SVIs * 2 parameters + 7 insensitive SVIs * 13 sensitive SVIs * 2 parameters) spectral-temporal features were generated as pairwise combinations of dynamics parameters obtained from sensitive and insensitive SVI. These features were then used for classification and regression, as described below.

#### 3.4.2 Binary classification into healthy and diseased canopies using spectral-temporal features

A PLSDA model using 4 components achieved a classification accuracy acc_int_ = 1.00, suggesting correct classification of each experimental plot as healthy or diseased canopy based on spectral-temporal features. In the external validation, the model achieved acc_gen_ = 0.86, thus correctly classifying 304/353 plots as healthy. This is slightly less accurate than the best time-point specific models (Figure 6).

#### 3.4.3 Regression models to quantify disease severity using spectral-temporal features

Overall, disease severity predictions from spectral-temporal features were similarly accurate as those obtained from time-point specific models based on reflectance spectra. The lowest RMSE was achieved when no control plots were used as training data using the PLS algorithm (RMSE = 0.068, *R*^2^ = 0.71). Differences in performance among algorithms were smaller than in time-point specific analyses, but the random forest performed relatively poorly also in this case (RMSE = 0.076, *R*^2^ = 0.62). Both tree-based models, and particularly the random forest, produced systematically biased predictions, which was not observable for PLS and ridge regression. In contrast to the time-point specific analyses, validation on the largely disease-free plots of the GABI panel suggested the necessity to include the control plots in the training data in order to obtain accurate predictions for low levels of disease, except for ridge regression. When these were included, tree-based models produced good estimates of the low disease-levels, whereas PLS and ridge regression still predicted disease severity of > 0.05 in a significant number of plots. The inclusion of the control plots resulted in a lower systematic bias of the tree-based predictions, while only marginally decreasing model performance (RMSE_adj_ = 0.074 and RMSE_adj_ = 0.076 for cubist and random forest, respectively; Figure 9A). Importantly, the inclusion of the control plots also strongly reduced or eliminated spatial patterns in predictions of the GABI panel, except for ridge regression (Figure 9B). Thus, cubist regression seemed to perform best when taking all evaluated aspects of model performance into account.

**Figure 9.**
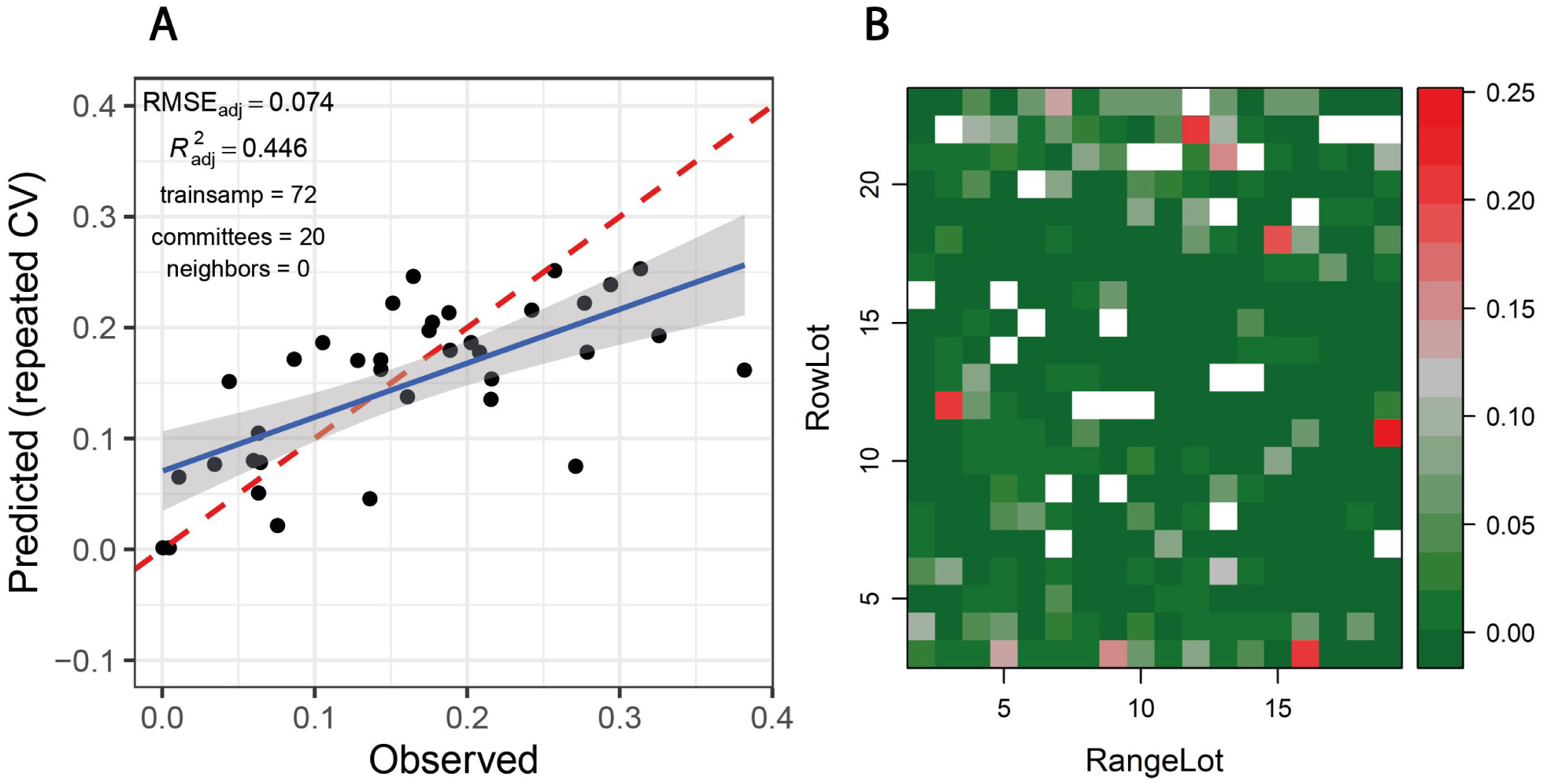
**(A)** Predicted vs. observed septoria tritici blotch (STB) severity levels of 36 artificially inoculated wheat plots. STB severity was measured on flag leaves using a combination of visual incidence scorings and scans of flag leaves exhibiting disease symptoms. Predictions were obtained from a cubist regression model based on spectral-temporal features for the same 36 plots and 36 non-inoculated control plots sown with the same genotypes. The broken red line represents the 1:1 line, the blue line represents the least squares line of the linear regression of predicted vs. observed values, the gray area represents the 95% confidence interval of the least squares line. **(B)** Spatial distribution of predicted STB severity levels of ~360 largely disease-free plots of the GABI wheat panel, grown next to the plots used as training dataset. White fields correspond to the plots contained in the training dataset.

#### 3.4.4 Feature selection

Feature selection was performed to identify the most important spectral-temporal features and to estimate the benefit of adding additional features. The difference between *M* derived from the modified chlorophyll absorption ratio index (MCARI2) and the structure insensitive pigment index (SIPI) was consistently the most informative spectral-temporal feature (Table S1). The MCARI2 was designed to estimate green leaf area index in crop canopies, whereas the SIPI measures pigment concentrations and ratios in leaves. This feature was retained as the last in all 30 resamples by the random forest and in 29 resamples by cubist. Following features had much increased ranks. The most influential features were all based on the *M* parameter of the Gompertz model, whilst other parameters were clearly less important. In particular, differences in change parameters did not seem to be informative of disease severity. Most selected features used the SIPI, R780/R740 and PRInorm indices as STB-insensitive index, even though they seemed to be somewhat more affected by the presence of disease than the FII on average (Table 3). There was little evidence for the existence of complementary information among the spectral-temporal features, as model performance was affected little by the sequential removal of features (Figure 10). However, the small sample size resulted in very high variance of the performance estimates obtained from the test set and contrasting patterns between the cross-validated training and the test performance estimates (Figure 10).

**Figure 10.**
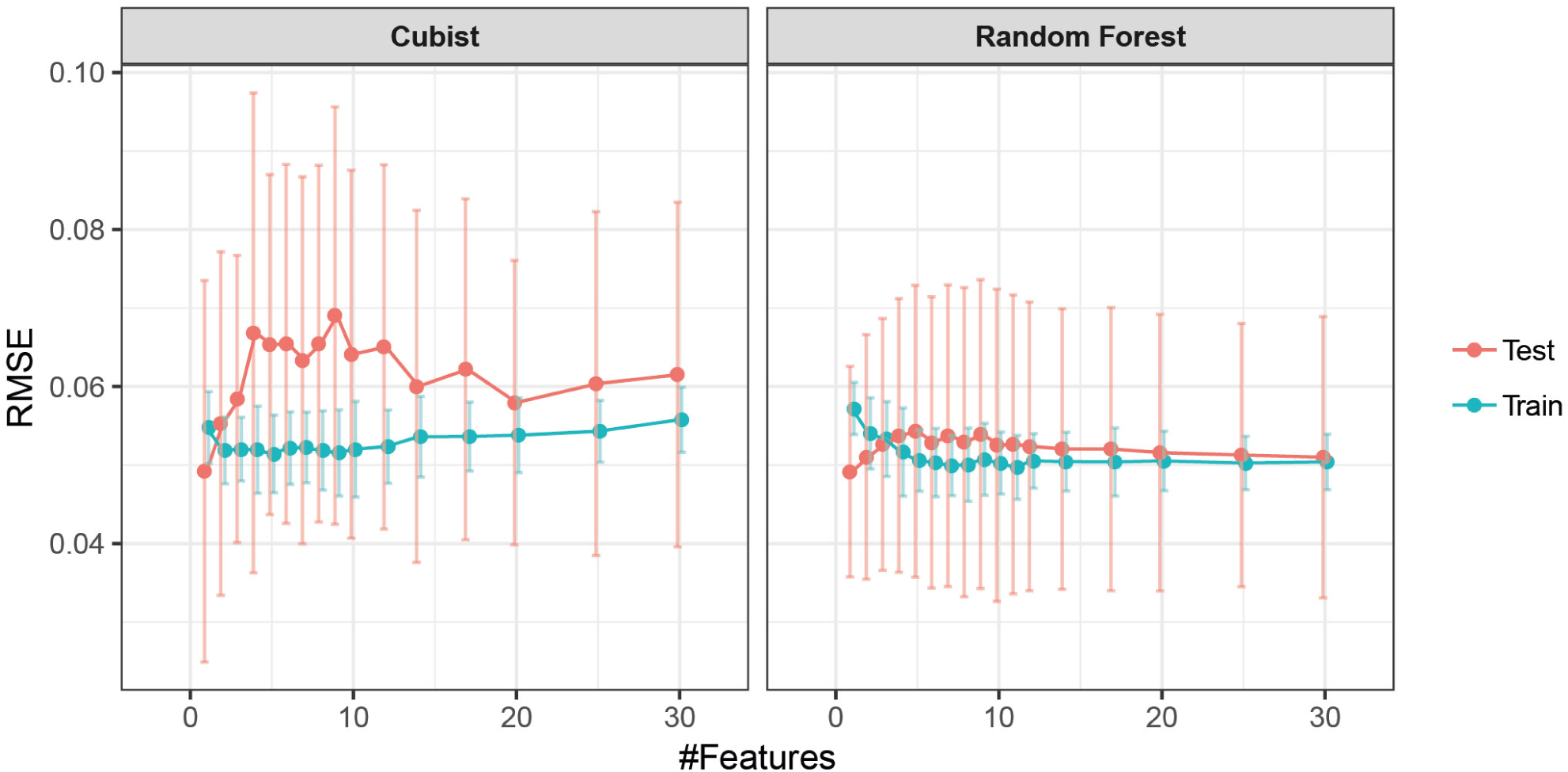
Performance profile of models based on spectral-temporal features to predict STB severity as a function of the number of spectral-temporal features used. Mean performance and standard deviation are shown based on 30 resamples of the data.

**Table 3.** Spectral vegetation indices (SVI) identified to be insensitive in their temporal dynamics to the presence or absence of septoria tritici blotch (STB) disease. For each dynamics parameter, a subset of eight SVI with the smallest difference (diff, GDD) or ratio (dimensionless) of the parameters between treatments was selected. The reported SVI were retained after subset selection based on pairwise correlations. The mean pairwise correlation (corr) is reported per dynamics parameter. Values in brackets report the minimum and maximum pairwise correlation.

| SVI | DynPar | diff/ratio | Pairwise corr |
| --- | --- | --- | --- |
| R780/R740 | t85 | 75.3 | 0.66 (0.26, 0.95) |
| CHLRE | t85 | 55.6 |  |
| DCNI | t85 | 45.6 |  |
| FII | t85 | 33.2 |  |
| GNDVI | t85 | 81.3 |  |
| PRInorm | t85 | 58.2 |  |
| SIPI | t85 | 45.3 |  |
| R780/R740 | M | 37 | 0.82 (0.60, 0.95) |
| CHLRE | M | 55.6 |  |
| DCNI | M | 25.7 |  |
| FII | M | 4.4 |  |
| GNDVI | M | 40.5 |  |
| Prinorm | M | 21.6 |  |
| SIPI | M | 22.7 |  |
| R780/R750 | b | 1.03 | 0.73 (0.48, 0.88) |
| CHLRE | b | 0.98 |  |
| MTCI | b | 0.95 |  |
| SR | b | 0.99 |  |
| R780/R750 | dur | 1.03 | 0.75 (0.53, 0.88) |
| CHLRE | dur | 1 |  |
| MTCI | dur | 1.02 |  |
| SR | dur | 0.99 |  |

## 4 Discussion

### 4.1 Limitations of time-point specific analyses

Current reflectance-based approaches to high throughput phenotyping of crop diseases under field conditions suffer from a lack of specificity and from insufficient robustness to genotypic diversity and environmental variability (i.e. context specificity). This problem has previously been described in detail with regard to growth stages of the crop, different phases in the pathogenesis and the presence of other stresses (Devadas et al., 2015; Zhang et al., 2012; Zheng et al., 2019).

Our results prominently illustrate context-specificity of the relationship between spectral reflectance and disease. Firstly, variation on a specific date in potentially disease-sensitive spectral features, such as the SIPI (*see* Bajwa et al., 2017; Yu et al., 2018), is quickly overridden by variation caused by advancing phenology (Figure 4C), illustrating the difficulty in defining thresholds or calibration curves. Secondly, unstable VIP values of single wavelengths in PLSDA models, systematic shifts in VIP patterns (Figure 6) and limited model applicability over time, even for the plots contained in the training dataset (Figure 7), illustrate marked short-term changes in the relationship between STB and spectral reflectance. Thirdly, the decreased classification accuracy on independent test plots (Figure 6) indicates context-specificity related to the effect of genotypes and, possibly, field heterogeneity.

There was a short period during late stay-green when classification models were transferable between time-points to some extent (Figure 6). This can be explained by the relatively synchronized appearance of moderate to high levels of STB in front of the relatively stable background of a stay-green canopy. In this intermittent phase, the signal caused by STB is strong compared to the noise caused by genotypic diversity and in-field measurements (see also Figure 4A). Nonetheless, the regression model for STB-severity based on reflectance spectra is still context-specific, as reflectance in the NIR (used as predictors) gradually decreases during the stay-green and senescence phase irrespective of the presence of STB (Figure 4A). NIR reflectance is also strongly affected by genotype morphology, canopy 3-D structure and canopy cover (Gutierrez et al., 2015; Jacquemoud et al., 2009) and is therefore not specific to STB if analyzed on a particular point in time. In addition, time-point specific models highlight the potential of detecting STB in different phases, using different spectral features. This potential would be left unused if only a short period would be targeted.

### 4.2 Potential of temporal changes in reflectance to detect and quantify STB

Due to the strong limitations of models based on reflectance spectra, we evaluated the potential of exploiting temporal changes in reflectance for disease detection and quantification instead. Models based on spectral-temporal features were characterized by a somewhat lower performance compared with models trained on reflectance spectra of a specific time-point (Figure 7A, Figure 8A). Nevertheless, classification accuracies were similar to the time-point specific PLSDA models, and regression models suggested that spectral-temporal features were also informative of disease severity. This is encouraging, particularly given the strongly contrasting morphological, canopy structural and stay-green properties of the genotypes comprised in the experiment.

### 4.3 Selected spectral indices and resulting spectral-temporal features

Even though high levels of STB developed during the stay-green phase in most artificially inoculated plots (Figure 4), several SVIs could be identified which displayed similar temporal patterns across treatments (Table 3). In particular, the flowering intensity index (FII; Stuckens et al., 2011), i.e. the normalized difference of R475 and R365, was found to be almost unaffected by STB (Figure 2A). In a previous study, we found that early physiological senescence of wheat canopies results in only (proportionally) small increases in reflectance at wavelengths shorter than 500 nm (unpublished data). Strong increases were observable only towards later stages of senescence. The observed insensitivity of the FII to STB likely results from the fact that STB affects only parts of the vegetation, initially mostly lower leaf layers, while significant amounts of healthy green tissue remain. Thus, FII values should change significantly only with the onset of rapid apical senescence, encompassing a generalized loss of green leaf area. It has been suggested that STB does not accelerate or anticipate apical senescence under a range of environmental conditions (Bancal et al., 2016). This is in line with the observed insensitivity of the FII to STB. Interestingly, the dynamic pattern of the structure insensitive pigment index (SIPI; Penuelas et al., 1995) was also found to be highly insensitive to STB. In contrast, this SVI was previously suggested as a potential surrogate for crop disease under field conditions (Bajwa et al., 2017; Yu et al., 2018). This Index was developed at the leaf scale to maximize sensitivity to the ratio between carotenoid and chlorophyll *a* concentrations (Car/Chl *a* ratio), while minimizing the effect of leaf surface and mesophyll structure (Penuelas et al., 1995). Provided the principles underlying the SIPI hold also for canopy level reflectance, a low sensitivity of the dynamic pattern to the presence of STB would be expected, as there seems to be no reason to expect a significant change at canopy level of the Car/Chl *a* ratio due to STB. STB causes the appearance of localized necrotic lesions; however, a general increase in the Car/Chl *a* ratio is not expected, unless STB accelerates or anticipates apical senescence, which does not seem to be the case (Bancal et al., 2016). The PRInorm was also among the most STB-insensitive SVIs. This SVI is based on the photochemical reflectance index (PRI), initially employed to measure changes in the relative levels of pigments in the xanthophyll cycle (Gamon et al., 1992). Over larger temporal scales, the PRI was shown to be strongly responsive to the Car/Chl ratio (Sims and Gamon, 2002). Zarco-Tejada et al. (2013) modified this SVI to decrease the effect of reduced canopy leaf area resulting from water stress. Thus, its insensitivity to STB can probably be explained in an analogous manner as for the SIPI.

The temporal patterns of water-sensitive SVIs such as the water index (WI; Peñuelas and Inoue, 1999) and the normalized difference water index (NDWI; Gao, 1996) were found to be highly sensitive to STB. Similarly, the disease water stress index (DSWI; Apan et al., 2004), which uses information from the water-sensitive SWIR and the NIR, was strongly affected in its temporal dynamics. In particular, water sensitive SVIs decreased much earlier for inoculated than for control plots, and the decrease occurred more gradually than in healthy plots (data not shown). This is in line with findings by Yu et al. (2018), who reported both the WI and NDWI to discriminate best between STB-diseased and healthy canopies in early stages of disease development. Several SVIs using reflectance in the RE and NIR also showed strongly contrasting temporal patterns (e.g. DSWI, NDVI, PSRI, and VI700). Similar to the SIPI, the plant senescence reflectance index (PSRI; Merzlyak et al., 1999) is highly sensitive to the Car/Chl ratio at the leaf level. However, NIR reflectance is used to normalize the difference between R677 and R500. It seems highly questionable whether the PSRI is particularly sensitive to the Car/Chl ratio in diverse germplasm at the canopy level. Variation in the PSRI seems to arise primarily from differences in canopy structure among genotypes and from canopy structural changes over time (Anderegg et al., 2019, *submitted*). The modified chlorophyll absorption ratio index (MCARI2; Haboudane et al., 2004), sensitive to green leaf area index, also showed strongly contrasting dynamic patterns (Figure 2B). The healthy-index (HI; Mahlein et al., 2013) was developed to separate healthy sugar beet leaf tissue from tissues affected by various foliar diseases. The prominent use of the RE by this index suggests that in our case, HI values are mostly driven by canopy structure and to a lesser extent chlorophyll absorption. Overall, our results thus suggest that SVIs sensitive to leaf internal structure and canopy structure are strongly affected by the presence of STB. This effect has been previously described for various patho-systems (*e.g.* Yu et al., 2018; Zhang et al., 2012; Zheng et al., 2019).

In summary, it seems that many of the derived spectral-temporal features can be interpreted as robust measures of STB-induced temporal changes in leaf internal structure, canopy structural parameters and canopy water content. These are obtained by normalizing the temporal dynamics of corresponding SVIs *via* the estimation of temporal changes in pigment ratios and reflectance at short wavelengths centered around 465 nm, likely representing physiological apical senescence. Thus, spectral-temporal features seem to well represent our hypothesis H_2_, as STB-affected plant and canopy traits are expressed relative to phenology-related traits.

### 4.4 Robustness of spectral-temporal features

As far as quantifiable in the framework of this experiment, models based on spectral-temporal features were robust to variation in genotype morphology, phenology, canopy cover and canopy 3D-structure as well as genotype-specific temporal changes thereof. Accurate predictions were obtained also for >300 diverse genotypes comprised in the GABI panel and grown in a large field experiment, suggesting robustness of the method to varying growing conditions arising from field heterogeneity (Figure 9B).

Results from feature selection suggested that a single spectral-temporal feature (i.e. the difference between *M* derived from MCARI2 and SIPI), relating structural changes in leaves and canopies to senescence-induced changes in pigment composition, was sufficient to achieve the performance illustrated above. Other stresses occurring during grain filling such as terminal drought stress and nitrogen shortages are likely to result in a similar decrease in green leaf area index. However, these stresses are also known to accelerate physiological senescence (Bogard et al., 2011; Distelfeld et al., 2014; Martre et al., 2006). Therefore, we speculate that the developed models may be moderately robust against the effect of common other stresses despite their simplicity. Yet, we conclude that our hypothesis H_3_ (i.e. that the combination of several spectral-temporal features representing the unique sequence and dynamics of separate events during pathogenesis could increase the specificity of the method) remains to be confirmed in larger experiments including other stress factors.

### 4.5 Multiple spectral vegetation indices to exploit temporal dynamics in reflectance

A key component of the proposed approach consists in summarizing hyperspectral data in terms of multiple SVIs and modelling of their temporal dynamics. Though this may result in the loss of some relevant information contained in reflectance spectra (Pauli et al., 2016), the use of SVIs presented a number of advantages here: (i) noise in temporal patterns was much reduced compared to reflectance values at single wavelengths, facilitating the fitting of parametric models; (ii) the inevitable subset selection step preceding feature combination could be based on objective criteria related to the form and purpose of SVIs; (iii) many of the used SVIs were designed specifically to maximize responsiveness to certain vegetation properties while minimizing the effect of common confounding factors, which is likely to also increase the robustness of derived spectral-temporal features (*see e.g.* Haboudane et al., 2004; Penuelas et al., 1995); and finally (iv) the procedure results in a summary of the hyperspectral dataset that is interpretable in terms of plant physiology and canopy characteristics, which also holds true for derived spectral-temporal features. Fitting parametric models to scaled SVI values may smooth out measurement errors related to single measurement dates, resulting for example from varying sun angle at measurement or short-term variation in illumination conditions. Thus, scaling SVI values and modelling their temporal dynamics reduces the effect of confounding factors on initial reflectance spectra and minimizes the effect of errors related to single measurements in the series.

### 4.6 Context and scope

In this study, we used a non-imaging spectroradiometer and manual feature engineering for disease detection and quantification. A high spatial resolution of imaging sensors has been deemed critical for disease detection, identification and quantification by others (Mahlein, 2016; Mahlein et al., 2012, 2010). The high potential of 2-D information in combination with deep learning methods for disease identification has been demonstrated recently (Fuentes et al., 2017; Mohanty et al., 2016). However, changes in spectral reflectance over time have also been shown to be highly informative at the leaf level (Mahlein et al., 2010; Wahabzada et al., 2015). To make use of the spatial and temporal dimensions under field conditions, individual lesions would arguably have to be tracked across time. Some solutions to this problem have been presented for close-range hyperspectral measurements (Behmann et al., 2018). However, similar solutions at the canopy level may be technically extremely challenging to implement and require extensive studies due to problems in tracking individual pixels or organs over time and in obtaining a clean spectral signal from objects with varying orientation. Existing approaches to make use of spectral, spatial and temporal information rely on automated and data-driven extraction of characteristic spectral features for diseased plants under controlled conditions (Thomas et al., 2018; Wahabzada et al., 2016, 2015). Here, promising results were achieved using a non-imaging sensor and manual feature extraction. This highlights that an improved understanding of potential confounding factors arising under field conditions may equally boost the potential of remote sensing methods for applications in crop breeding.

We developed and validated the presented method to facilitate robust in-field detection and quantification of STB. However, the underlying concepts should be transferrable to different problems, such as the detection and quantification of other foliar diseases. Several features of the proposed approach (e.g. exploiting plot-based relative changes in reflectance over time, combining sensitive and insensitive features, or the SVI-based parameterization of temporal dynamics) may also be valuable in quantifying other breeding-relevant traits, such as the timing and dynamics of nitrogen remobilization.

## 5 Conclusion

Here, we tested the possibility to detect and quantify STB relying exclusively on relative changes in spectral reflectance over time, which is expected to minimize confounding effects on spectral reflectance arising from genotypic diversity and environmental conditions. Our results demonstrated the feasibility of the proposed approach and suggested that resulting models were robust against variation in several common nuisance factors. Specifically, it appears that the temporal dynamics in green leaf area index when set in relation to the dynamics of physiological apical senescence is highly indicative of the presence of STB infections and of STB severity. Time-resolved measurements of the MCARI2 and the SIPI spectral vegetation indices could allow to assess these traits at very high throughput, facilitating time-resolved large-scale screenings of breeding nurseries.

Larger calibration experiments will offer the opportunity to evaluate the inclusion of additional spectral-temporal features that better capture relevant information in different phases of pathogenesis. This is likely to improve sensitivity and specificity of resulting models, which should also be tested in more detail. Furthermore, the evaluation of the scalability to unmanned aerial vehicles will represent a crucial step towards application of such methods in breeding programs.

## 7 Author Contributions

All authors planned the experiment. AH created the experimental design. PK prepared and carried out pathogen inoculation. AM and PK designed the STB assessment strategy and supervised STB scorings, collections and scanning of leaves. JA performed reflectance measurements. JA performed leaf scan analysis, conceived the proposed data analysis approach, analyzed the data and drafted the manuscript. AM and AH supervised the project. All authors contributed to, read and approved the final version of the manuscript.

## 9 Acknowledgments

We thank Prof. Dr. Achim Walter for input to the overall conception of the study and Mr. Hansueli Zellweger for field management and crop husbandry. The authors gratefully acknowledge fruitful discussions with Mr. Philipp Baumann and Dr. Helge Aasen. Pathogen inoculation was supported by Mr. Reto Zihlmann. Measurements in the field and STB assessment campaigns were supported by Ms. Kelbet Nagymetova, Ms. Delphine Piccot, Mr. Michel Colombo and Mr. Pablo Bovy. The project was partially funded by the Swiss Federal Office of Agriculture (FOAG). AM and PK gratefully acknowledge financial support by the Swiss National Science Foundation through Ambizione grant PZ00P3_161453.

## 10 Data Availability Statement

The raw data supporting the conclusions of this manuscript will be made available by the authors, without undue reservation, to any qualified researcher.

